# Pericyte-Like Cells Undergo Transcriptional Reprogramming and Distinct Functional Adaptations in Acute Lung Injury

**DOI:** 10.1101/843805

**Authors:** CF Hung, S Holton, YH Chow, WC Liles, SA Gharib, WA Altemeier

**Author notes:** Address correspondence to: Chi F Hung, MD, Division of Pulmonary and Critical Care, University of Washington, Center for Lung Biology, Box 358052, 850 Republican St, Seattle, WA 98109.

## Abstract

**Background:** We previously reported on the role of pericyte-like cells as functional sentinel immune cells in lung injury. However, much about the biological role of pericytes in lung injury remains unknown. Lung pericyte-like cells are well-positioned to sense disruption to the epithelial barrier and coordinate local inflammatory responses due to their anatomic niche within the alveoli. In this report, we characterized transcriptional responses and functional changes in pericyte-like cells following activation by alveolar components from injured and uninjured lungs in a mouse model of acute lung injury (ALI).

**Methods:** We purified pericyte-like cells from lung digests using PDGFRβ as a selection marker and expanded them in culture as previously described (1). We induced sterile acute lung injury in mice with recombinant human Fas ligand (rhFasL) instillation followed by mechanical ventilation (1). We then collected bronchoalveolar lavage fluid (BALF) from injured and uninjured mice. Purified pericyte-like cells in culture were exposed to growth media only (control), BALF from uninjured mice, and BALF from injured mice for 6 and 24 h. RNA collected from these treatment conditions were processed for RNAseq. Targets of interest identified by pathway analysis were validated using *in vitro* and *in vivo* assays.

**Results:** We observed robust global transcriptional changes in pericyte-like cells following treatment with uninjured and injured BALF at 6 h, but this response persisted for 24 h only after exposure to injured BALF. Functional enrichment analysis of pericytes treated with injured BALF revealed activation of immuno-inflammatory, cell migration and angiogenesis-related pathways, whereas processes associated with tissue development and remodeling were down-regulated. We validated select targets in the inflammatory, angiogenesis-related, and cell migratory pathways using functional biological assays *in vitro* and *in vivo*.

**Conclusion:** Lung pericyte-like cells are highly responsive to alveolar compartment content from both uninjured and injured lungs, but injured BALF elicits a more sustained response. The inflammatory, angiogenic, and migratory changes exhibited by activated pericyte-like cells underscore the phenotypic plasticity of these specialized stromal cells in the setting of acute lung injury.

## Introduction

Pericytes are specialized perivascular mural cells that are in direct contact with the endothelium in the microvasculature. They play a critical mechanistic role in angiogenesis during development (2, 3). However, the biology of lung pericytes in the post-developmental state, both in homeostasis and in response to physiologic stress, remains poorly characterized. Published studies on pericytes have relied on a combination of molecular markers and histology to define the cellular population studied. Although these methods greatly enrich the population of cells under investigation for pericytes, the cellular populations are invariably heterogeneous. We, therefore, refer to these cells as “pericyte-like” cells.

Reports on pericyte-like cells have shown that they may assume different biological functions in homeostasis and in response to injury. During homeostasis, pericyte-like cells contribute to endothelial integrity in the central nervous system (4). Following organ injury, however, pericytes may exhibit functional changes in response to injury. Several reports suggest pericyte-like cells are important progenitors of activated myofibroblasts in organ fibrosis (5, 6). In animal models of organ injury, pericyte-like cells have been shown to participate in local inflammatory responses through elaboration of chemokines and upregulation of adhesion molecules that interact with immune cells (7–9).

We have previously described lung pericyte-like cells as functional sentinel immune cells in the murine lung (1). Similar to findings in other organs, lung pericyte-like cells, selected by platelet-derived growth factor receptor beta (PDGFRβ) expression, respond to a number of toll-like receptor (TLR) agonists *in vitro* by elaborating cytokine and adhesion molecule expression. Furthermore, ablation of pericyte-like cells prior to induction of bleomycin-induced lung injury resulted in attenuation of inflammatory cells and total protein in the alveolar compartment, suggesting that these cells contribute to the inflammatory response in the lung (10). Thus, emerging evidence suggests lung pericyte-like cells undergo significant changes that may be functionally relevant to inflammation and subsequent repair in the lung.

Although we previously showed that lung pericyte-like cells exhibit pro-inflammatory properties during lung injury, the full extent of functional changes remains incompletely characterized. To address this gap in our understanding of the biology of pericyte-like cells during injury, we used transcriptomics to characterize the spectrum of functional changes in activated pericyte-like cells. We further demonstrated the functional upregulation of select pathways identified by transcriptomic analysis.

## Results

### Transcriptional response in pericyte-like cells

Using RNA-seq, we first interrogated global transcriptional changes in pericyte-like cells exposed to uninjured BALF and to injured BALF compared to control conditions (media only) at 6 and 24 h (Fig 1A). In pericyte-like cells exposed to uninjured BALF for 6 h, we observed 2104 differentially regulated genes (FDR < 0.01) compared to media only control. Of these 2104 genes, 1249 were up-regulated and 855 were down-regulated. For the 24 h exposure, however, only 106 genes remained differentially regulated compared to media only control (68 up-regulated and 38 down-regulated genes).

**Fig 1.**
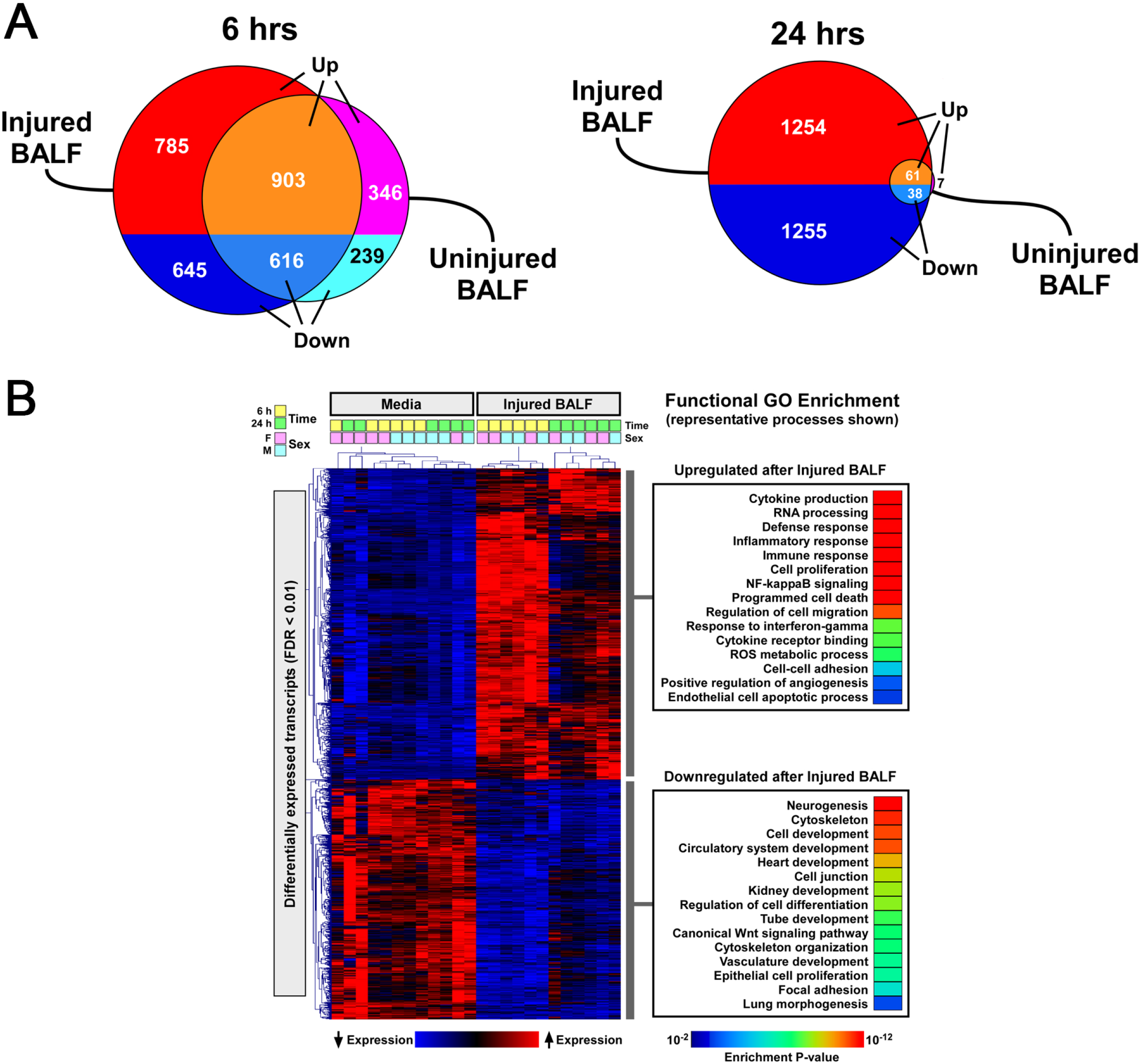
(A) Diagrams illustrating the number of differentially expressed genes in pericyte-like cells exposed to uninjured BALF and injured BALF compared to media-only control (FDR<0.01). Diagram on the left illustrates the number of differentially expressed genes at 6 hours. Diagram on the right illustrates the differentially expressed genes at 24 hours. In sharp contrast to 6 h time point, only a small number of genes (99 genes) remain differentially regulated at 24 hours in pericytes treated with uninjured BALF whereas the changes persisted in pericytes exposed to injured BALF. (B) Functional GO enrichment analysis of differentially expressed genes in pericyte-like cells exposed to injured BALF compared to media-only control. Representative processes that are upregulated and downregulated are shown on the right.

In pericyte-like cells exposed to injured BALF for 6 h, we found 2949 differentially regulated genes compared to the media-only control condition. Of these genes, 1688 were up-regulated and 1261 were down-regulated. In sharp contrast to the uninjured BALF treatment, a vast number of differentially regulated genes persisted at 24 h in pericyte-like cells treated with injured BALF: we observed 2615 differentially expressed genes compared to media only control after 24 h of exposure (1322 up-regulated and 1293 down-regulated genes).

These results indicate that alveolar components in homeostatic conditions can induce short-term global transcriptional changes in pericyte-like cells even in the absence of alveolar damage. However, only contents from injured alveolar space cause sustained transcriptional responses in pericytes in contrast to stimulation with uninjured BALF.

To further characterize the transcriptional response of pericyte-like cells exposed to injured BALF, we conducted functional GO analysis on the subset of genes persistently differentially expressed at 24 h (Fig 1B). Consistent with our previous findings, immune and inflammatory responses were significantly activated in pericyte-like cells exposed to injured BALF (1). Interestingly, processes associated with regulation of cell migration and angiogenesis were also up-regulated. In contrast, pathways associated with remodeling and development were down-regulated in pericyte-like cells exposed to injured BALF. Results from the functional enrichment analysis indicated that pericyte-like cells undergo widespread transcriptional reprograming in the presence of inflammatory stimuli. These findings imply that beyond immuno-inflammatory roles, pericyte-like cells exhibit additional phenotypic changes during acute lung injury that alter their function within the microvascular niche.

### Validation of select genes in functionally enriched GO categories

We selected several representative genes mapping to distinct enriched GO functional categories for validation (Fig 2). Changes in the expression of these candidate genes were examined at the transcript level by qPCR. Early transcriptional changes in pro-inflammatory genes *Cxcl1*, *Il6*, and *Ifng* were observed in pericyte-like cells treated with uninjured and injured BALF for 6 h. Treatment with uninjured BALF led to increased expression of *Cxcl1*, *Il6*, *and Ifng* compared to media only control. At 24 h, treatment with uninjured BALF continued to induce the expression of *Cxcl1*. Treatment with injured BALF continued to induce the expression of *Cxcl1* and *Ifng*. In contrast to pro-inflammatory genes, expression of selected angiogenic genes such as *Vcam1*, *Nos2*, and *Angptl4* was only induced by injured BALF but not by uninjured BALF.

**Fig 2.**
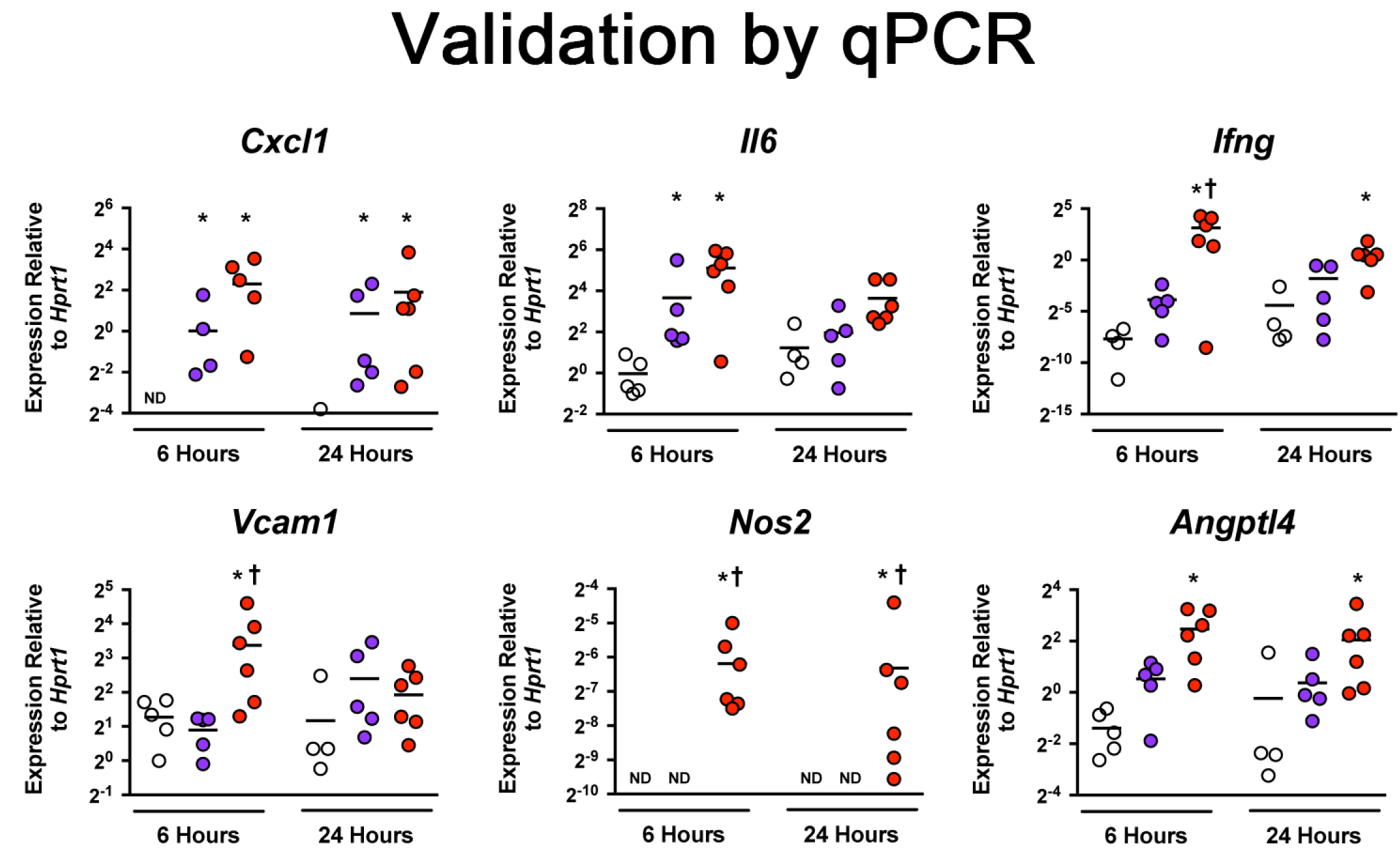
Confirmation of differentially expressed genes by qPCR. Cultured pericyte-like cells were exposed to media-only control (white), uninjured BALF (blue), or injured BALF (red) for 6 and 24 hours. Select genes from inflammatory and angiogenic pathways are presented. Level of mRNA expression is expressed relative to *Hprt* gene expression. (Bar=mean, n≥4, *p<0.05 vs. media-only control, †p<0.05 vs uninjured BALF)

### Validation by protein assays

Elevated levels of immune-related and angiogenesis-related cytokines identified by RNA-seq were detected in cell culture supernatants and cell lysates of uninjured BALF and injured BALF treated pericyte-like cells. Cell culture supernatants from pericyte-like cells after 24 h of treatment with uninjured BALF showed significantly elevated levels of IL-6 and CCL20 by ELISA compared to media-only controls (Fig 3A). Cell culture supernatant from pericyte-like cells treated with injured BALF showed elevated levels of CXCL1, CXCL2, IL-6, and CCL20 compared to media-only controls (Fig 3A). Overall, treatment of pericyte-like cells with both uninjured and injured BALF induced a proinflammatory phenotype in pericyte-like cells.

**Fig 3.**
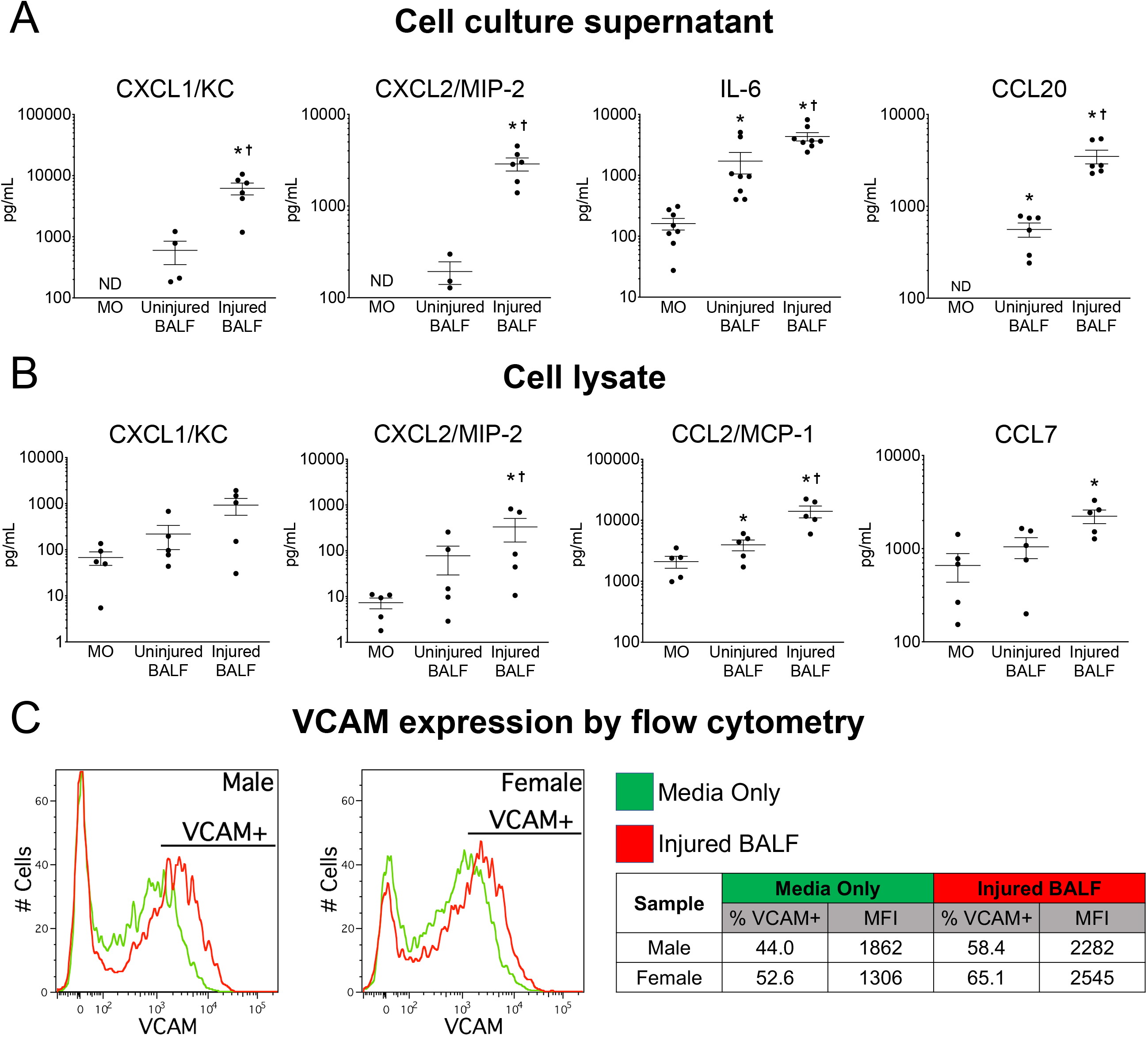
(A) Concentration of selected proinflammatory cytokines in cell culture supernatants after 24 hours of treatment in media-only control (MO), Uninjured BALF, and Injured BALF. (B) Concentration of selected proinflammatory cytokines in cell lysates from pericyte-like cells after 24 hours of treatment in media-only control (MO), Uninjured BALF, and Injured BALF. (Bar=mean±SEM, n=5, *p<0.05 vs. MO, ^†^p<0.05 vs uninjured BALF) (C) Histograms for VCAM immunostaining in pericyte-like cells treated with media-only control (green line) and Injured BALF (red line). Gating for VCAM+ population is shown in the histogram. Table shows % population that is VCAM+ and the mean fluorescence intensity (MFI) of each sample.

Because detection of cytokines in cell culture supernatant may reflect the sum of pericyte-like cell-derived cytokines and cytokines already present in the BALF, select cytokines were evaluated by ELISA using cell lysates obtained from BALF stimulated cells to further confirm the pro-inflammatory effect of BALF on pericyte-like cells. Cells treated with uninjured BALF demonstrated increased CCL2 in cell lysates at 24 h compared to media-only control (Fig 3B). Treatment with injured BALF further augmented the pro-inflammatory profile with elevated levels of CXCL2, CCL2, and CCL7 in cell lysates compared to media-only control (Fig 3B).

We previously demonstrated that pericytes upregulate ICAM-1 expression *in vivo* when challenged with lipopolysaccharide (LPS) by oropharyngeal aspiration (1). In this study, RNA-seq data also revealed upregulation of VCAM-1, which was confirmed at the mRNA level (Fig 2). We further assessed the expression of VCAM-1 in pericyte-like cells at the protein level by flow cytometry. We isolated pericyte-like cells from a male and a female mouse. Cells treated with injured BALF for 24 h showed elevated expression of VCAM-1 compared to media-only controls (Fig 3C).

### Demonstration of upregulation of Angpt4 in vivo

One of the most highly upregulated genes in injured BALF-exposed pericyte-like cells identified by RNA-seq is *Angptl4*. Through *in situ* RNA hybridization (RNAScope^®^), we demonstrated the upregulation of angiopoietin L4 transcript (*Angptl4*) in pericyte-like cells *in vivo* following sterile lung injury (Fig 4). We observed Tdtomato mRNA labeling in perivascular stromal cells (pericyte-like cells) in PDGFRβ-CreER^T2^;Ai14 mice. In the absence of lung injury, we observed very low levels of *Angptl4* expression localized to Tdtomato-labeled cells (Fig 4B). *Angptl4* expression was significantly induced in Tdtomato-labeled cells with lung injury.

**Fig 4.**
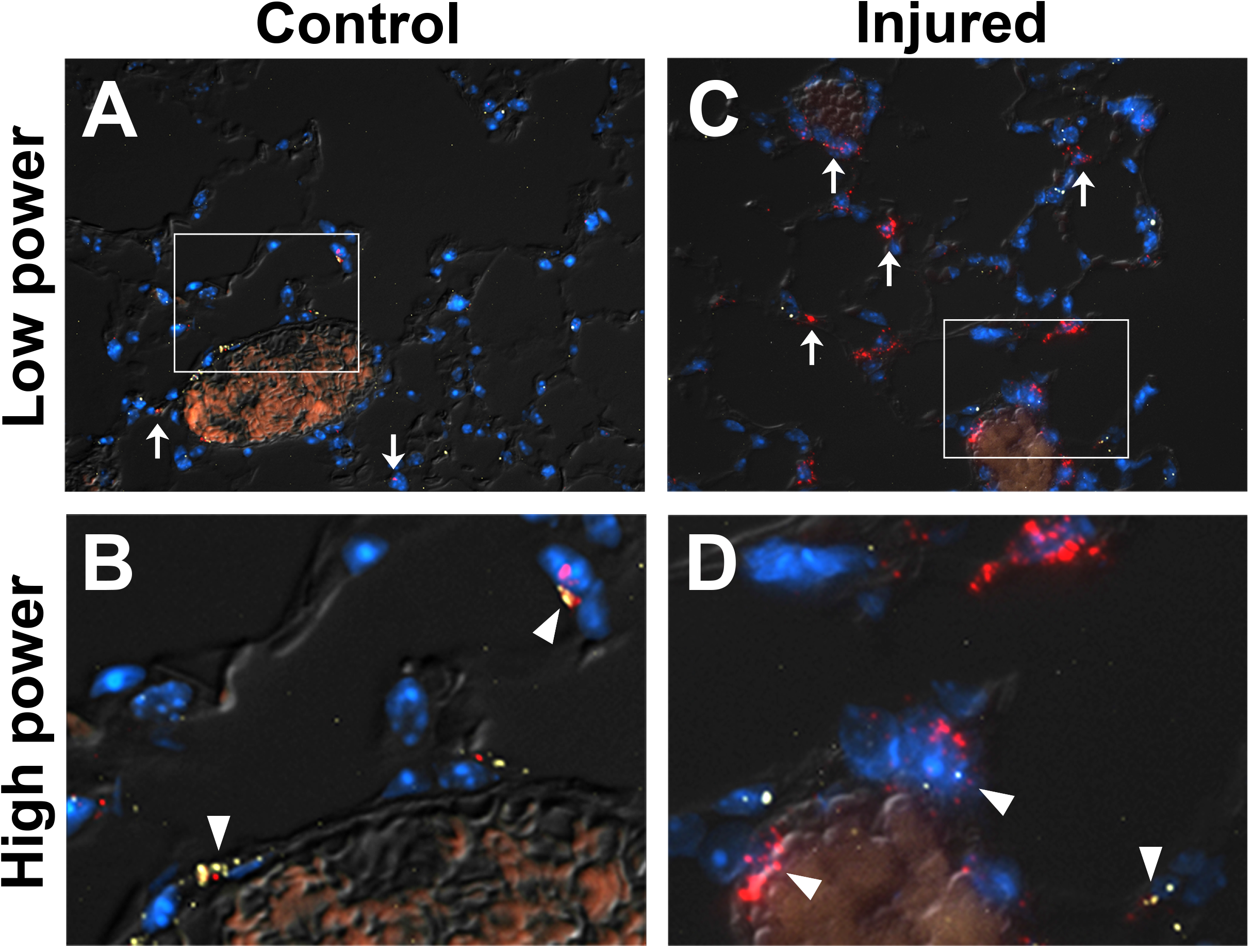
RNAScope^®^ In Situ Hybridization of *Angptl4* mRNA (red) and *Tdtomato* mRNA (yellow) in PDGFRβ-CreER^T2^;Ai14 mouse lung sections. Nuclei were counterstained in DAPI (blue) and images were superimposed on DIC images. (A) Image of uninjured lung section at 40X. Examples of *Angptl4* mRNA expression are highlighted by arrows (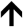). (B) Expanded view of inset in (A). Cells with co-expression of *Angptl4* mRNA (red) and *Tdtomato* mRNA (yellow) are highlighted by arrowheads (▴). (C) 40X image of injured mouse lung. Arrows (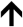) indicate examples of *Angptl4* mRNA expression (red). (D) Expanded view of image inset in (C). Arrowheads (▴) indicate examples of cells with co-expression of *Angptl4* mRNA (red) and *Tdtomato* mRNA (yellow).

### Enhanced matrix invasion by activated pericyte-like cells

We observed enrichment in pathways associated with regulation of cell migration. To understand whether these transcriptional responses translated to functional changes, we examined matrix invasiveness in pericyte-like cells stimulated with uninjured and injured BALF. Compared to media-only controls, uninjured and injured BALF induced significant increases in Matrigel invasiveness in pericyte-like cells (Fig 5A and 5B). We further examined whether the observed findings could be attributed to differences in proliferation between treatment groups, and we did not find a significant difference in 24 h proliferation between media-only and BALF-exposed groups (Fig 5C).

**Fig 5.**
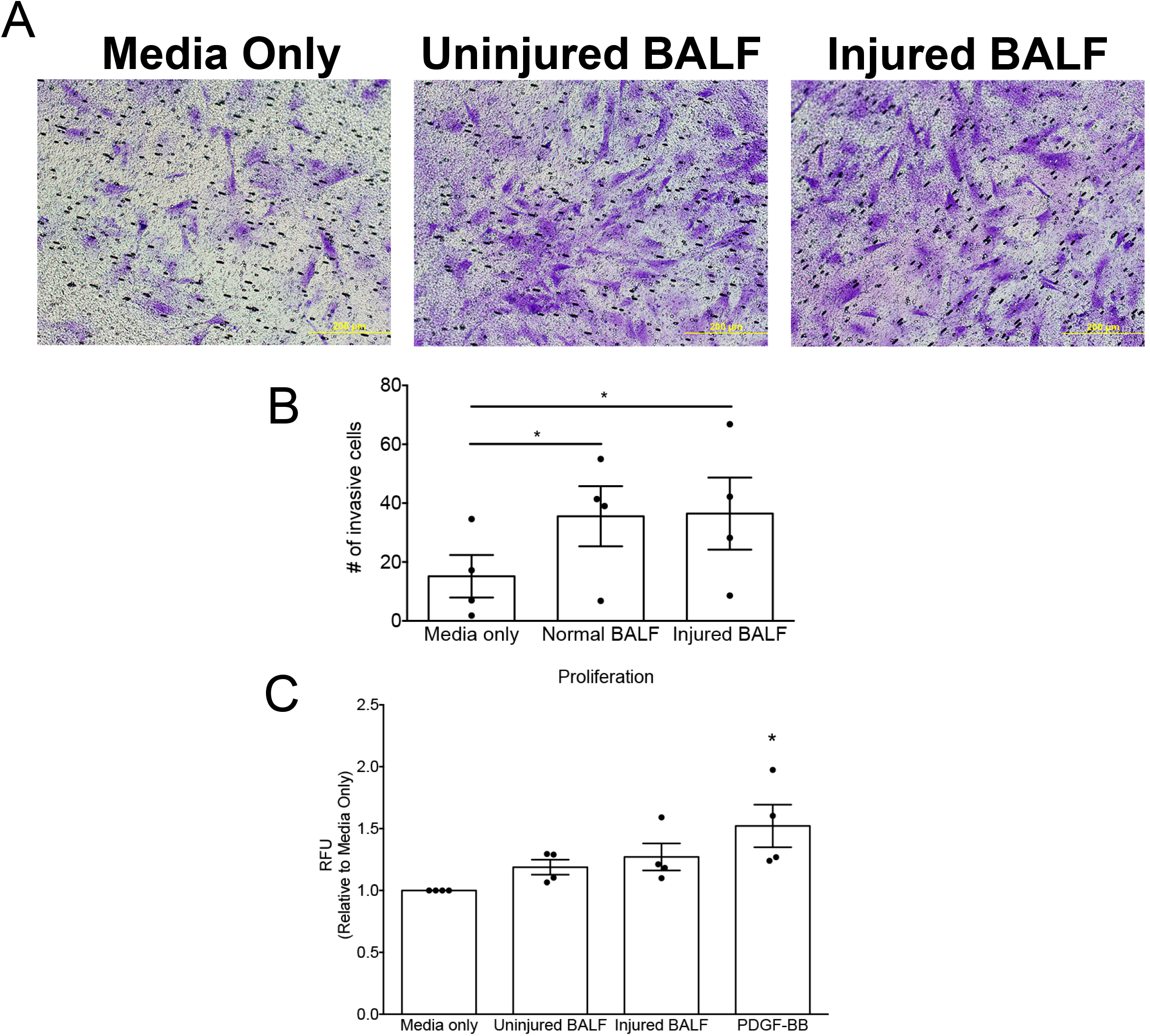
(A) Representative images of pericyte-like cells (purple) that invaded across Matrigel-coated membranes under different treatment conditions at 24 hours (bar = 200μCm). (B) Number of invaded cells at 24 hours were counted over 5 high power fields. (Bar graphs represent mean±SEM, n=4, *p<0.05 vs. media-only control.) (C) Proliferation assay at 24 hours measured by fluorescence units. Proliferation under the indicated treatment conditions is normalized to media-only control and exp

## Discussion

Large gaps in knowledge continue to exist in our understanding of how lung pericyte-like cells respond to physiologic stress. Accumulating evidence suggests pericyte-like cells exhibit a degree of functional plasticity in response to organ injury. Our group previously reported evidence demonstrating the relevance of pericyte-like cells in both acute lung injury and in fibrosis using animal models (1, 6, 10). During lung homeostasis, an intact alveolar epithelium protects pericyte-like cells from alveolar contents. However, in the setting of acute lung injury, pericyte-like cells are exposed to a complex microenvironment characterized by loss of intact epithelial barrier and infiltration of alveolar components in the lung interstitium. These components may consist of native alveolar contents, necrotic debris, damage-associated molecular patterns, pathogens, pathogen-associated molecular patterns, and inflammatory cytokines. How pericyte-like cells behave in this environment of alveolar injury remains incompletely understood.

In this study, we characterized the spectrum of injury response in pericyte-like cells through transcriptional profiling. Consistent with our previous report, we observed significant upregulation in inflammation response pathways when lung pericyte-like cells were exposed to uninjured and injured BALF (1). The inflammatory response was much more robust and sustained in pericyte-like cells exposed to injured BALF. However, even in the absence of injury, alveolar components from uninjured BALF were able to induce transient transcriptional changes in pericyte-like cells, supporting the notion that pericyte-like cells are primed to detect breakdown in the alveolar epithelial barrier. We verified these changes in selected targets at transcript and protein levels. Importantly, these changes were also observed in pericyte-like cells *in vivo*, suggesting a functional relevance for transcriptomic results.

Interestingly, we also observed enrichment in other pathways such as cell migration and regulation of angiogenesis through gene set enrichment analysis. These changes in pericyte-like cells in response to acute lung injury were previously unrecognized and hint at the possibility that pericyte-like cells play an active role in its microvascular niche following injury. The ability for pericyte-like cells to migrate within and from their native perivascular niche and disrupt endothelial barrier function may hold important clues about their functional roles in acute lung injury and subsequent repair. Evidence for such an active role has been demonstrated in other organs. For example, in one study of pericytes in cremaster muscle, the authors showed that pericytes upregulated leukocyte adhesion molecules and reorganized their cellular orientation within the basement membrane to guide neutrophils in the subendothelium (7). This type of cellular trafficking requires coordinated changes that include breakdown of extracellular matrix, enhanced mobility of pericyte-like cells within the perivascular niche, and upregulation of leukocyte adhesion molecules. Results from our transcriptional profiling and *in vitro* functional studies of activated lung pericyte-like cells suggest these processes may also have relevance in lung injury. Whether these functional changes orchestrate or augment the trafficking of immune cells in the lung warrants additional study.

Pericytes play an important role in angiogenesis during embryonic and early development. However, little is known about their biologic activity during homeostasis and after organ injury in adults. Our functional GO analysis revealed angiogenic-regulation as one of the persistently upregulated pathways in pericyte-like cells exposed to injured BALF for 6 and 24 hours. Within this pathway, *Angptl4* was one of the most highly upregulated genes and we demonstrated its upregulation *in vivo* following lung injury. Angiopoietin-like 4 (ANGPTL4) has two biologically active domains following proteolytic cleavage. Evidence suggests N-terminal domain is involved in lipoprotein metabolism (11). The C-terminal region (cANGPTL4) is secreted and evidence in tumor biology suggests it may mediate vascular leak (12). An examination of human lung tissue samples from the 1918 and 2009 influenza pandemics revealed that angiopoietin-like 4 was one of the most highly upregulated genes in the lung (13). Moreover, published reports suggest antagonism of cANGPTL4 in pulmonary infection models ameliorated pulmonary vascular leak and damage (14, 15). In our study, we observed significant upregulation of *Angptl4* in pericyte-like cells both *ex vivo* and *in vivo* following stimulation. Given the physical proximity of pericytes and endothelial cells, the ANGPTL4 signaling axis may be an important pathway in how pericytes regulate endothelial integrity during acute lung injury.

An important limitation in our study is that the transcriptomic profile of activated pericyte-like cells was obtained *ex vivo*. The microenvironment in the microvascular niche is complex and the transcriptomic response of pericyte-like cells *in vivo* may differ from our findings. Therefore, we validated select pathways and genes of interest *in vivo*. Interrogation of other pathways of interest identified through transcriptomics will similarly require validation in animal models to ensure biological relevance.

In summary, we demonstrate that lung pericyte-like cells undergo robust transcriptional reprogramming upon exposure to alveolar components from uninjured, and most notably, from injured BALF. The transcriptomic approach proved to be a helpful hypothesis-generating tool that facilitated the identification of novel functional pathways in activated pericyte-like cells that were previously understudied. In addition to their inflammatory roles, pericyte-like cells may alter their interactions with the endothelium and the extracellular matrix during acute lung injury. These findings highlight the phenotypic plasticity of pericyte-like cells under physiologic stress. Further characterization of these functional changes during injury may provide deeper insights into the mechanisms of lung injury and novel cellular targets to modify pathologic inflammation.

## Methods

### Animals and reagents

The University of Washington Institutional Animal Care and Use Committee approved all experiments involving the use of mice described in this report.

Antibodies used for magnetic-activated cell sorting (MACS) included biotin-conjugated anti-CD31 (clone 390), anti-CD45 (clone 30-F11), anti-CD326 (clone G8.8), and PE-conjugated anti- PDGFRβ (clone APB5) (eBioscience). Magnetic columns and antibody-conjugated magnetic beads were purchased from Miltenyi Biotec (San Diego, CA). Liberase TL Research Grade enzyme mix and DNase I used in lung digestion were purchased from Roche Applied Science (Pleasanton, CA). Luminex® Multiplex and ELISA assays for the detection of chemokines and cytokines were purchased from R&D Systems (Minneapolis, MN).

Recombinant human Fas ligand (rhFasL) was purchased from Enzo Life Sciences (Farmingdale, NY). Recombinant IL-1β was purchased from PeproTech (Rocky Hill, NJ).

Primer probe sets for qPCR were purchased from Integrated DNA Technologies (IDT, Coralville, IA).

### Acute lung injury model

We induced sterile acute lung injury in mice by using a model that combines rhFasL and mechanical ventilation as previously described (1). Briefly, male and female mice 8-12 weeks of age were given rhFasL by oropharyngeal aspiration to induce alveolar epithelial injury. To augment injury, we mechanically ventilated the mice 24 hours following FasL administration for a duration of 6 hours at a tidal volume of 8 cc/kg. Following mechanical ventilation, mice were euthanized and their trachea were exposed. A small incision was made through which a metal catheter was inserted and secured to the trachea by sutures. To collect bronchoalveolar lavage fluid (BALF), we inflated the lungs with 1ml sterile phosphate buffered saline (PBS) followed by aspiration and collection of the lavage. BALF from either injured mice (we will refer to this as “injured BALF” hereafter) or uninjured littermates (“uninjured BALF”) were collected. BALF from mice was further processed for tissue culture by collecting supernatants from centrifugated samples and passing the supernatants through 0.22 micron sterilizing filters.

### Pericyte isolation and culture

Single cell preparations from mouse lung digests were prepared as previously described (1). Briefly, single cell preparations were expanded in culture in 0.2% gelatin-coated T75 flasks for 3-5 days until confluence. Cultured cells then underwent negative selection using CD31, CD45, and CD326 to isolate the stromal cell fraction (unlabeled cells). Pericyte-like cells were selected from stromal cells using PDGFRβ as positive selection marker. Selected cells were cultured on gelatin-coated tissue culture flasks in cell culture media (DMEM/F12 supplemented with 10% FBS, 1% ITS, and 1% penicillin/streptomycin) until use in experiments.

### Transcriptional analyses

#### RNA collection

Cultured pericyte-like cells were seeded at a density of 25,000 cells/well in a 12-well tissue culture plate or 250,000 cells/well in a 6-well tissue culture plate. Following attachment, they were exposed to culture media with reduced serum (DMEM/F12 supplemented with 2.5% FBS, 1% ITS, and 1% penicillin/streptomycin) mixed 1:1 with sterile PBS (“media only” hereafter), a 1:1 mixture of reduced serum media and uninjured BALF, or a 1:1 mixture of reduced serum media and injured BALF. At 6 h and 24 h following exposure to treatment conditions, cell culture supernatants and cell lysates were collected for further processing and analysis. RNA was collected from treated cells using RNeasy Mini Kit per manufacturer’s protocol (QIAGEN). RNA integrity was confirmed with a Bioanalyzer 2100 (Agilent, Santa Clara, CA). RNA for RNA sequencing and for qPCR validation studies were collected from independent and separate experiments.

#### RNA sequencing

RNA-seq was performed at University of Washington’s Northwest Genomics Center (http://nwgc.gs.washington.edu/). Briefly, library generation was performed using TruSeq Stranded mRNA library prep kit (Illumina, San Diego, CA) using 1 μg total RNA. Sequencing was performed using an Illumina HiSeq 4000 instrument using paired end 75 bp reads. Base calls were generated in real-time on the HiSeq instrument, and RTA-generated BCL files were converted to FASTQ files. Sequence read, base quality and other QC metrics were computed and checked with RNASeQC (v2.6.4) (16). Sequences were aligned to mouse genome using the gencode (M11) annotation and STAR (v2.5.2b) on mouse GRCm38.primary_assembly (17). Transcript abundances were quantified using RSEM (v1.2.31) (18).

#### RNA-seq data analysis

After filtering low count transcripts, differentially expressed genes between conditions (injured BALF vs. media control, uninjured BALF vs. media control) at each time point (6 h, 24 h) were identified using “DESeq2” (19) package within the R statistical environment. Multiple hypothesis testing correction was implemented using Benjamini-Hochberg’s false discovery rate (FDR) analysis, with an FDR < 0.01 designating significant differential gene expression (19). Functional enrichment analysis was performed on differentially expressed genes using the online application Webgestalt (20) based on Gene Ontology (GO) annotations and hypergeometric statistical modeling to determine enrichment *P*-values. Multiple hypothesis testing adjustment was performed using Benjamini-Hochberg’s method and an FDR < 0.01 was used to identify significantly enriched GO categories. All RNA-seq data meeting MINSEQE (Minimum Information About a Next-generation Sequencing Experiment) have been deposited at Gene Expression Omnibus repository.

### Confirmatory studies

#### Validation by qPCR

Pericyte-like cells were cultured as previously described. They were treated with media only, BALF from uninjured animals, or BALF from injured animals. RNA for qPCR validation studies was collected from independent experiments separate from RNAseq experiments. RNA was isolated per protocol as described above. cDNA was synthesized using M-MLV Reverse Transcriptase (Invitrogen, ThermoScientific) in a 150ng/20 μl reaction using a BioRad thermocycler. The cDNA was then diluted 1:5 and qRT-PCR reactions were set up in triplicate using primers synthesized from IDT and SensiMix SYBR & Fluorescin kit (Bioline, Meridian Bioscience). qRT-PCR was done using a Corbett Research RG 6000 thermocycler. A comparative CT method (2^−ΔΔCt^) was used to obtain expression fold change relative to housekeeping gene *hrpt*.

#### Flow cytometry

We used flow cytometry to assess protein expression changes in pericyte-like cells under the same treatment conditions as described above. Both cell-surface and intracellular targets were interrogated. To enhance detection of intracellular targets, we added a commercially available protein transport inhibitor (GolgiStopTM, BD Biosciences) to the culture media 2 h before harvest. Cultured cells were collected after treating with Accutase. Collected cells were treated with Fc Block (BD Biosciences) followed by immunostaining with specific antibodies conjugated to fluorophores. Immunostaining of intracellular targets was performed using Perm/Fix system (BD Biosciences) to fix and permeabilize cells prior to intracellular immunostaining. Fixable viability dye eFluor450 was used to identify live cells.

#### Cytokine detection

We added GolgiStopTM (BD Biosciences) to culture media 2 h before harvest to enhance detection of normally secreted cytokines in cell lysates. At the indicated timepoints, cell culture supernatants and cell lysates were collected. Treated cells were lysed in protein lysis buffer with protease and phosphatase inhibitors (Roche). Selected cytokine targets were measured in cell culture supernatants and cell lysates with commercially available Luminex® Multiplex Assays (R&D Systems) per manufacturer’s protocol. Analytes that fell below the level of detection were assigned a value equal to one-half the lower limit of quantification of the assay for the purposes of statistical analysis. In graphical presentation of the data, only detectable samples with detectable levels of analytes are represented with data points. In cases where none of the biological samples in a treatment group had measurable levels, the data is presented as none detected (ND).

#### *In situ* RNA analysis of lung sections

PDGFRβ-CreER^T2^ transgenic mice expressing a tamoxifen-inducible Cre recombinase under the endogenous PDGFRβ promoter (B6.Cg-Tg(Pdgfrb-cre/ERT2)6096Rha/J, The Jackson Laboratory) were crossed with Tdtomato reporter mice Ai14 (B6.Cg-*Gt(ROSA)26Sor*^*tm14(CAG-tdTomato)Hze*^/J, The Jackson Laboratory) to generate the bitransgenic reporter mice PDGFRβ-CreER^T2^;Ai14. Reporter mice were fed a tamoxifen containing diet postnatally between weeks 3-7. Cells expressing PDGFRβ during tamoxifen exposure and their daughter cells were identified by permanent expression of the red fluorescent protein Tdtomato. Tamoxifen-exposed PDGFRβ-CreER^T2^;Ai14 mice underwent lung injury by rhFasL/mechanical ventilation as described above. Harvested lungs were inflated with 10% normal buffered formalin (Sigma) at 25cm H_2_O column pressure, fixed in 10% normal buffered formalin overnight and stored in 70% EtOH until processing for paraffin embedding.

Fluorescently labeled RNA probes against *Angptl4* and *Tdtomato* were purchased from Advanced Cell Diagnostics, Inc. Hybridization of RNA targets in paraffin-embedded lung sections using the RNAScope^®^2.0 Assay (Advanced Cell Diagnostics, Inc.) was performed per manufacturer’s protocol (21).

### Cell-culture assays

#### Matrix invasion assay

CytoSelect^TM^ 24-well Cell Invasion Assay (Cell Biolabs, Cat # CBA-110) was used to assess matrix invasion. We conducted the experiment per manufacturer protocol with the following specifications: Insert membranes were pre-wetted with serum-free DMEM/F12 for 10 minutes at room temperature. Following the step, pericyte media were added to the well (lower chamber). We seeded the inserts (upper chamber) with 2×10^5^ cells in pericyte media. The cells were allowed to attach to the membrane at 37C, after which the pericyte media were removed from the lower and upper chambers. Cell culture media corresponding to the experimental conditions were then placed in the lower and upper chambers, and the cells were allowed to incubate for 24 hours until fixation and cell counting per manufacturer protocol.

#### Cell proliferation assay

Cell proliferation was evaluated using CyQuant Cell Proliferation Kit (ThermoFisher, #C7026). Briefly, pericyte-like cells were seeded at a density of 1×10^4^ cells per well in 96-well tissue culture-treated black plates with clear bottom for fluorescence applications (Corning Cat. #3904). Cells were allowed to attach for 3 hours in pericyte medium, after which the medium was replaced with treatment media. Cells were incubated with treatment media for 24 hrs, gently washed in sterile PBS once, and then frozen in −80C. After thawing, proliferation assay was performed per manufacturer’s protocol. Fluorescence was detected using Synergy 4 microplate reader (Biotek). Proliferation is reported as relative fluorescence units compared to treatment control (media only).

#### Statistical analysis

For qPCR and cytokine quantification in BALF and cell lysates, data were log-transformed prior to statistical analysis. One-way ANOVA with Tukey’s correction was used when comparing 3 or more experimental groups. All tests were performed in Prism 8 (GraphPad). *P* values of less than 0.05 were considered statistically significant.

## Author Contributions

CFH, SAG, WCL, and WAA were involved in research design; CFH, SH, and YHC were involved in acquiring data; CFH, SH, SAG, WCL, and WAA were involved in analyzing data; WAA provided reagents; CFH, SH, SAG, WCL, and WAA were involved in writing of this manuscript

## Acknowledgements

Funding for this work is provided by grants from the NIH: R01HL122895 (WAA) and K08HL127075 (CFH)

